# Severe distortion in the representation of foveal visual image locations in short term memory

**DOI:** 10.1101/2021.10.03.462912

**Authors:** Konstantin F. Willeke, Araceli R. Cardenas, Joachim Bellet, Ziad M. Hafed

**Author notes:** **Correspondence to** Ziad M. Hafed, Werner Reichardt Centre for Integrative Neuroscience, Tübingen University, Otfried-Müller Str. 25, Tübingen, 72076, Germany.

## Abstract

The foveal visual image region provides the human visual system with the highest acuity. However, it is unclear whether such a high fidelity representational advantage is maintained when foveal image locations are committed to short term memory. Here we describe a paradoxically large distortion in foveal target location recall by humans. We briefly presented small, but high contrast, points of light at eccentricities ranging from 0.1 to 12 deg, while subjects maintained their line of sight on a stable target. After a brief memory period, the subjects indicated the remembered target locations via computer controlled cursors. The biggest localization errors, in terms of both directional deviations and amplitude percentage overshoots or undershoots, occurred for the most foveal targets, and such distortions were still present, albeit with qualitatively different patterns, when subjects shifted their gaze to indicate the remembered target locations. Foveal visual images are severely distorted in short term memory.

## Introduction

The representation of visual spatial locations in short term memory has been one of the most classic means for studying the neural mechanisms of cognitive control in modern-day systems neuroscience (Bruce & Goldberg, 1985; Funahashi, Bruce, & Goldman-Rakic, 1989; Gnadt & Andersen, 1988; Gnadt, Bracewell, & Andersen, 1991; Hikosaka & Wurtz, 1983; Kojima, 1980). Experimentally, reporting a remembered target location with a saccadic eye movement has proven extremely useful in demonstrating how different cortical and subcortical brain regions may maintain a memory trace of visual targets (Constantinidis & Procyk, 2004), and it has also been equally important for oculomotor control studies in dissociating sensory and motor responses (Edelman & Goldberg, 2001; Mays & Sparks, 1980; Mohler & Wurtz, 1976; Willeke et al., 2019).

One mechanism for short term memory maintenance in multiple brain areas is the persistence of neural activity associated with remembered stimulus locations, even in the absence of sensory drive (Constantinidis & Procyk, 2004; Wimmer, Nykamp, Constantinidis, & Compte, 2014). Such persistence, with temporal drift, can help explain a variety of distortions in memory-based task performance, for example, as a function of how long a location needs to be maintained in short term memory (Wimmer et al., 2014). In addition, such persistence can reveal certain systematic biases with respect to whether stimuli occupy a single visual quadrant or multiple visual quadrants (Leavitt, Pieper, Sachs, & Martinez-Trujillo, 2018), which in turn enables linking visual field asymmetries in perception to representations of remembered target locations. For example, visual performance along the horizontal and vertical retinotopic meridians is different (Carrasco, Talgar, & Cameron, 2001; Liu, Heeger, & Carrasco, 2006; Talgar & Carrasco, 2002), and this difference is maintained in tasks involving short term memory (Montaser-Kouhsari & Carrasco, 2009). The fact that such visual meridian effects may be related to tissue magnification in visual cortical areas (Benson, Kupers, Babot, Carrasco, & Winawer, 2021; Silva et al., 2018; Van Essen, Newsome, & Maunsell, 1984) might then suggest that other known distortions in short term memory tasks, such as foveal biases in remembering peripheral target locations (Kerzel, 2002; Sheth & Shimojo, 2001), may also be related to how visual space is represented in topographic maps.

If remembering visual target locations depends on both how visual space is topographically represented as well as on how memory information is neurally maintained, then an important remaining open question is whether foveal visual locations are recalled veridically or not, given the normally very high acuity nature of foveal vision in humans. Here, we investigated this question and found that remembering a foveal visual location as near to the line of sight as 0.1 deg (6 min arc) is subject to severe distortion, even after very short memory delay intervals. This paradoxical foveal distortion in visual short term memory emerges with or without a constant landmark being available during the response phase of the task, and, perhaps most importantly, it is also qualitatively different depending on whether the remembered location is reported with an eye movement response or with another response modality.

## Results

We asked human subjects to fix their gaze on a small, central spot. During fixation, we briefly flashed, for 59 ms, a similar spot at a pseudorandom location between 6 min arc and 12 deg of eccentricity. After a memory interval of 300-1100 ms, the fixation spot disappeared, and a response cursor appeared in its place. The subjects moved the response cursor via a button box to the remembered location of the flash, and they were free to move their eyes during cursor control (Methods) (Willeke et al., 2019). We found a large distortion in the reported locations for targets presented at foveal visual eccentricities, very near to the line of sight. Such distortion also occurred: when we changed the response modality for indicating the remembered locations; when we maintained a central visual landmark during the subjects’ response execution; and also when we asked the subjects to indicate the remembered locations with eye movements.

### A strong diagonal bias and eccentricity overshoot in remembered foveal target locations

Figure 1A shows all possible target locations that we tested in our button cursor movement experiment (Methods). We densely sampled foveal eccentricities in all directions, and we also sampled extrafoveal locations. In the left two panels of Fig. 1B (labeled ‘Foveal targets’), we plotted, for two example foveal target locations (blue), a sampling of response indications by the subjects (faint red dots indicating individual trials). The average response location for each target, our variable of interest in this study (Methods), is indicated by a saturated red dot. For the purely diagonal target at an eccentricity <15 min arc (leftmost panel in Fig. 1B; ‘Diagonal’), the average memory-guided response was directionally accurate, but it exhibited an amplitude overshoot. For the second foveal target location at an angle of 30 deg below the right horizontal meridian (and still with an eccentricity <15 min arc), the average response not only overshot the target, but it was also strongly biased in direction towards the diagonal axis (middle panel; compare red and blue lines). This diagonal bias largely disappeared for the third example target location shown in Fig. 1B (rightmost panel); this target location still had a direction of 30 deg from the horizontal meridian, but it now was at a much larger extrafoveal eccentricity >5 deg (rightmost panel). Therefore, when remembering foveal visual target locations, our subjects exhibited a strong and diagonal bias for oblique locations, along with an eccentricity overshoot, which was much reduced at larger extrafoveal eccentricities.

**Figure 1.**
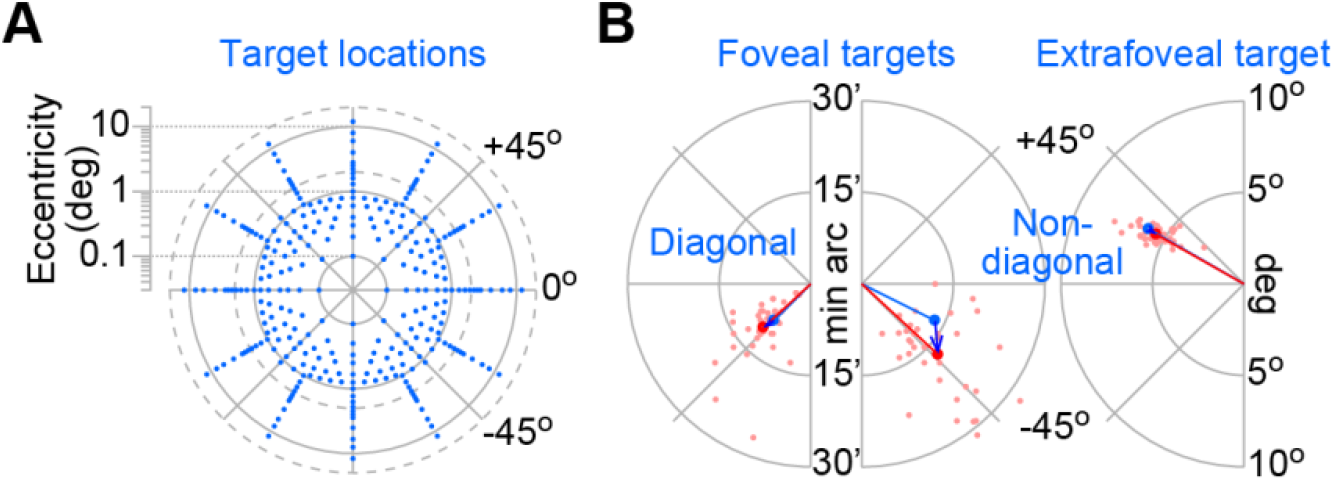
A strong distortion in remembered foveal target locations. **(A)** All target locations tested in our short term memory button cursor experiment. Each dot shows a potential remembered target location on a log-polar plot. In each trial, a brief flash appeared at one location. After a few hundred milliseconds, the display was blanked, and the subjects had to move a cursor to the flash’s remembered location. **(B)** Example response biases in remembered target locations. The left two panels show two example foveal targets at an eccentricity <15 min arc. Each target is shown as a blue dot (with a blue line connecting display center to the target). The faint red dots show example response locations. For the leftmost panel, the responses overshot the target, but their directions were (on average) correct (the red dot shows the average response location). For the middle panel, a similar foveal target eccentricity had a non-diagonal direction from the horizontal meridian; here, the responses were additionally strongly biased towards the diagonal axis. The blue arrow shows the mislocalization that took place (connecting target location to average response position). The rightmost panel shows an example non-diagonal target location (similar to the middle panel: 30 deg direction from the horizontal meridian), but now at a much larger eccentricity (note the different eccentricity scale). In this case, the responses were directionally-accurate, and there was no overshoot.

To visualize results across all target locations and eccentricities, we plotted, for each target, a vector connecting its true location and the average response location (similar to the blue arrow in the middle panel of Fig. 1B). The results, across data from all 7 subjects, are shown in Fig. 2A. Non-cardinal foveal target locations were strongly skewed towards the diagonal direction in each visual quadrant, and this happened at all tested foveal eccentricities (Fig. 2A). Note how there was always an amplitude overshoot, although such an overshoot was quite small for the target locations along the cardinal axes (purely horizontal and purely vertical directions). Moreover, the amplitude overshoot increased in size for the diagonal foveal targets, for which the direction errors were minimal (the response vectors for the purely diagonal targets were still largely diagonal, resulting in minimal direction errors). Therefore, complementary to known foveal biases in visual short term memory representations for peripheral, extrafoveal targets (Kerzel, 2002; Sheth & Shimojo, 2001), we saw strong overshoots in remembering target locations with small, foveal visual eccentricities. Most critically, we concurrently observed a severe diagonal bias in all non-cardinal foveal target locations.

**Figure 2.**
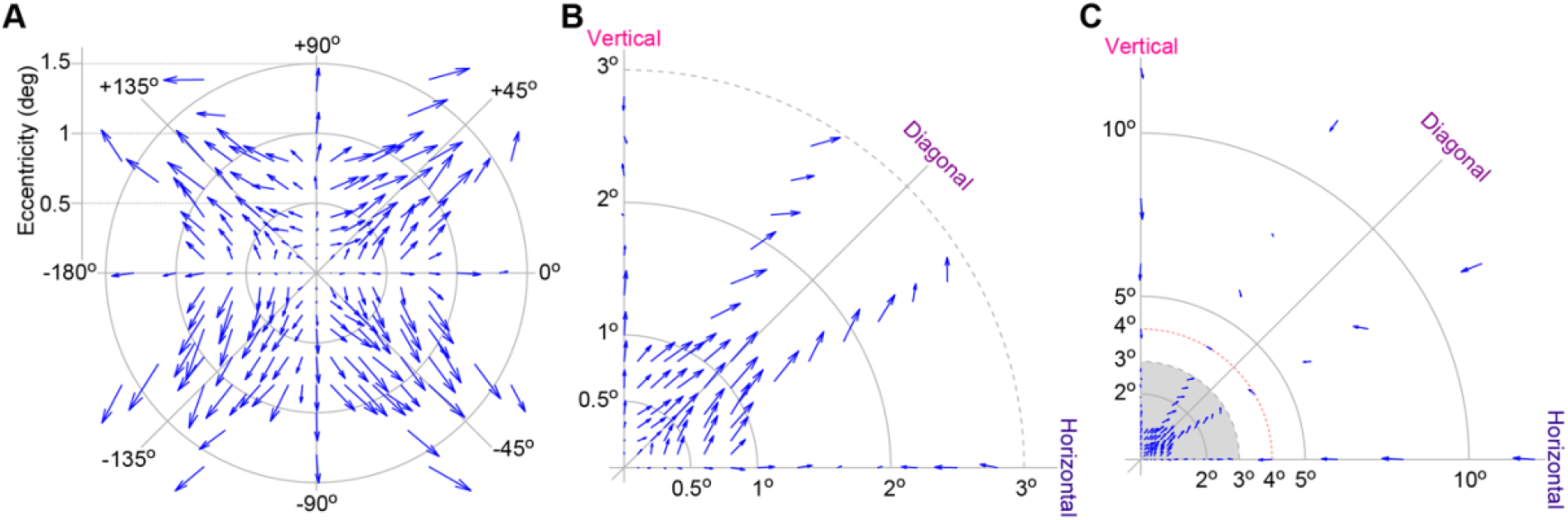
Distorted representation of foveal visual image locations in short term memory. **(A)** Systematic mislocalization of remembered foveal target locations as revealed by a quiver plot. The origin of each vector is the true target location, and the end of it (the arrowhead) is the average indicated location (similar to the blue vector shown in the middle panel of Fig. 1B). There was an amplitude overshoot for all tested foveal target locations (outward arrows from the origin in all directions). Moreover, there was a strong diagonal bias for non-cardinal target locations. Cardinal targets on the horizontal and vertical meridians were associated with the least errors. **(B)** Same data as in **A**, but after remapping all targets and responses to one quadrant (and after including parafoveal eccentricities). Note how the cardinal targets between 2 and 3 deg eccentricity were associated with a foveal bias (undershoots), but oblique targets at the same eccentricities were still associated with amplitude overshoots. **(C)** The gray region shows the data from **B**, and the rest of the figure shows results from more eccentric locations. The red dashed line shows that non-cardinal (oblique) targets flipped from overshoots to undershoots at an eccentricity of 4 deg. Figure S1 shows consistent results from individual subjects.

For further characterizing this strong diagonal bias, we exploited the general symmetry of the effect shown in Fig. 2A. We, therefore, remapped the target and response locations to one quadrant, for easier representation, and we plotted the memory-based mislocalizations. The results are shown in Fig. 2B. The strong diagonal bias at foveal eccentricities was robustly observed. Interestingly, a qualitative flip in memory-based localization error from being a response amplitude overshoot at foveal eccentricities to a response amplitude undershoot at higher eccentricities (the known foveal bias) occurred along the cardinal axes at an eccentricity of 2 deg (Fig. 2B); that is, overshoots in memory-based response localization occurred at less than 2 deg target eccentricities, whereas foveal biases (undershoots) occurred at more than 2 deg target eccentricities. However, non-cardinal locations still exhibited amplitude overshoots at eccentricities larger than 2 deg. For example, in Fig. 2B, eccentricities between 2 and 3 deg showed amplitude overshoots for non-cardinal remembered target locations, but they showed foveal biases (undershoots) at cardinal target locations. The flip from overshoots to foveal biases in memory-based target localizations, for non-cardinal target locations, occurred at an eccentricity of 4 deg (red dashed line in Fig. 2C). In terms of direction errors, the diagonal biases for non-cardinal target locations persisted even for eccentricities >10 deg, although they were milder than in the foveal visual representation (Fig. 2C; also see Fig. 3A).

**Figure 3.**
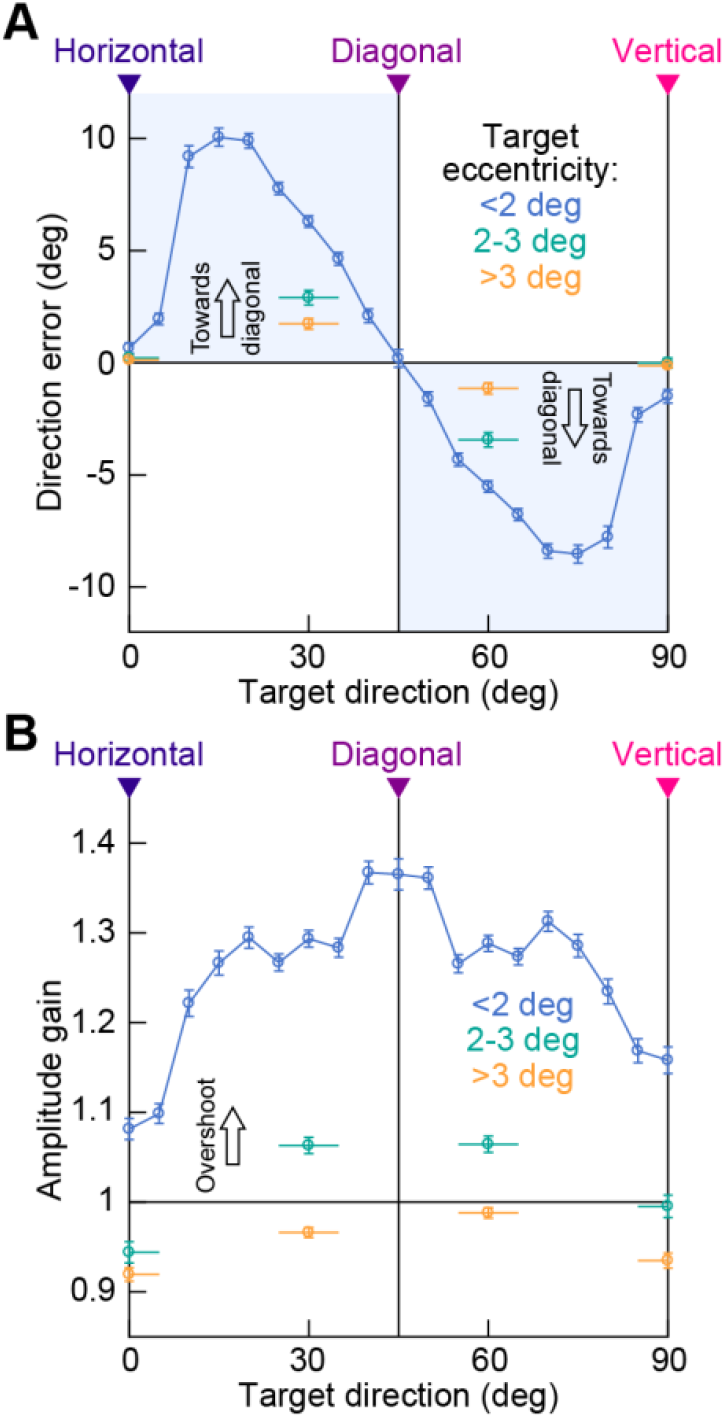
Direction and amplitude errors when remembering foveal and extrafoveal visual target locations. **(A)** For all remapped target and response locations (as in Fig. 2B, C), we plotted direction error as a function of target direction from the horizontal meridian. Direction error was the difference between the average response direction and true target direction. The blue curve shows results from foveal target eccentricities. Cardinal target directions were associated with minimal, but non-zero, direction errors. Non-cardinal target directions were associated with a strong diagonal bias, with near-horizontal targets having positive direction errors and near-vertical targets having negative direction errors (shaded regions). Direction error was practically zero at purely diagonal target locations. For extrafoveal target locations (green and orange), cardinal direction errors were essentially eliminated. Non-cardinal diagonal biases were much milder than in foveal eccentricities. **(B)** Same analyses but for response amplitude gain (ratio of average response eccentricity to true target eccentricity). Foveal target eccentricities (blue) exhibited strong overshoots, which peaked at purely diagonal directions. Eccentric targets (>3 deg) were associated with foveal biases (undershoots) (Kerzel, 2002; Sheth & Shimojo, 2001). Eccentricities between 2 and 3 deg (green) had mild overshoots for non-cardinal directions, as in Fig. 2. Error bars denote s.e.m. across all observations. Figure S1 shows the results from individual subjects, and Fig. S2A-D shows the effects of memory interval length on response distortions.

To further quantify the direction and amplitude errors in memory-based target localization, we binned target locations into different direction and amplitude bins. For foveal targets (<2 deg eccentricity), we created a running average of direction bins (15 deg bin widths and with running steps of 5 deg). In these bins, 0 deg direction would represent horizontal target locations and 90 deg direction would represent vertical target locations (as in the remapped representations of Fig. 2B, C). For each direction bin, we then calculated the average direction error between response direction and target direction (Methods). The resulting curve is shown in Fig. 3A (<2 deg target eccentricity). Other than for purely cardinal target locations, foveal targets close to the horizontal meridian (with directions less than 45 deg) were associated with positive direction errors (which systematically decreased in amplitude with increasing target direction), and foveal targets close to the vertical meridian (with directions larger than 45 deg) had negative direction errors. These results are consistent with an overall directional bias towards the diagonal axis (shaded regions in Fig. 3A; consistent with Fig. 2). For purely diagonal targets, the direction errors were minimal.

Interestingly, horizontal target locations (0 deg direction on the x-axis in Fig. 3A) were associated with lower absolute direction errors than vertical target locations (90 deg) (0.654 deg +/- 0.2305 deg s.e.m. versus 1.5 deg +/- 0.3043 deg s.e.m. direction error; an approximately 2.3 fold difference). This observation is reminiscent of better visual performance on the horizontal versus vertical visual meridian studied at larger peripheral eccentricities (Carrasco et al., 2001; Montaser-Kouhsari & Carrasco, 2009; Talgar & Carrasco, 2002), but here it was found in a memory-guided localization task and also extended to the foveal visual image region.

As for amplitude errors, we used the same directional binning as in Fig. 3A, and now measured the ratio of average response eccentricity to target eccentricity. For foveal targets (<2 deg target eccentricity), there was up to approximately 35% amplitude overshoot when remembering foveal visual target locations (Fig. 3B). The overshoots were largest for the purely diagonal targets (Fig. 3B), for which the direction errors were minimal (Fig. 3A), and the overshoots were the smallest for purely cardinal target locations. Moreover, overshoots for purely horizontal targets were less strong than overshoots for purely vertical targets (8.1% +/- 1.18% s.e.m. versus 15.8% +/- 1.48% s.e.m.; a 1.95 fold difference), again reminiscent (in terms of superior horizontal versus vertical meridian performance) of visual meridian effects reported at larger eccentricities (Carrasco et al., 2001; Montaser-Kouhsari & Carrasco, 2009; Talgar & Carrasco, 2002). All of these results are consistent with the observations summarized in Fig. 2.

The strong diagonal biases and amplitude overshoots that we observed were also most significant for foveal visual target locations. For larger remembered target eccentricities (extrafoveally), diagonal biases for oblique targets were still present in Fig. 3A (see the data points for 2-3 deg target eccentricities in green and >3 deg target eccentricities in orange), but they were much smaller in size than for foveal targets. For example, at 30 deg and 60 deg directions from the horizontal meridian, the absolute direction errors for targets >3 deg in eccentricity were 1.74 deg +/- 0.241 deg s.e.m. and 1.14 deg +/- 0.251 deg s.e.m., respectively, but they were 6.29 deg +/- 0.255 deg s.e.m. and 5.52 deg +/- 0.257 deg s.e.m. for targets <2 deg in eccentricity (Fig. 3A). This was a factor of 3.6-4.8 difference. Moreover, in terms of response amplitudes, targets at eccentricities >3 deg were consistently associated with known foveal biases (amplitude undershoots) (Kerzel, 2002; Sheth & Shimojo, 2001), which is the exact opposite of the overshoots that we observed for targets <2 deg in eccentricity.

Therefore, remembering recently presented visual target locations at foveal visual eccentricities is associated with strong distortions. These distortions are much larger in size, relative to the target eccentricities (and also opposite in sign in terms of amplitude errors), than distortions of more eccentric target location representations.

We also confirmed that the results in Figs. 2, 3 were robust at the individual subject level. For example, Fig. S1A-D shows representations of memory-based foveal target mislocalizations from 4 individual example subjects (in a format similar to that used in Fig. 2). Moreover, Fig. S1E, F shows all individual subject curves from the analyses of Fig. 3 for the foveal targets. All subjects showed a clear direction bias towards the diagonal axis, and they all showed amplitude overshoots, which were weakest for the purely cardinal target locations. Moreover, even though memory intervals in our task were relatively short (<1100 ms; Methods), we still observed an influence of memory interval length on the amplitude of the response distortion in our task (Fig. S2A-C), and this effect was not explained by a potential difference in response reaction times (Fig. S2D). Therefore, there is a large and robust distortion in the representation of the foveal visual image region in short term memory.

### Persistence of foveal image distortions in short term memory with a foveal visual landmark

In the above results, there was no visual landmark (other than the response cursor) during the final response phase of the task, and the subjects were also free to move their eyes during this phase. Moreover, cursor movement could take up to several seconds to reach the location intended by the subjects (Methods; Fig. S3A). We, therefore, asked whether the distortion that we observed so far (Figs. 1-3) was caused by any or all of the above factors. We conducted a second experiment, which only differed in the response modality. In this mouse pointing variant of the experiment, the trial ended with the persistence of the central fixation spot on the display, and the appearance of a computer mouse pointer (at a random location; see Methods). The subjects were instructed to maintain gaze on the central spot and use the computer mouse to point and click at the remembered target location. We used a similar range of foveal and extrafoveal target locations (Fig. 4A). Therefore, in this version of the task, we maintained gaze position near the original location (thus maintaining, on average, retinotopic correspondence during the visual presentation and response phases of the trials); we kept a foveal visual landmark on the display during the response phase, just like there was a foveal visual landmark at target flash onset; and, finally, we ensured a much faster reaction time than with cursor movement (Fig. S3B).

**Figure 4.**
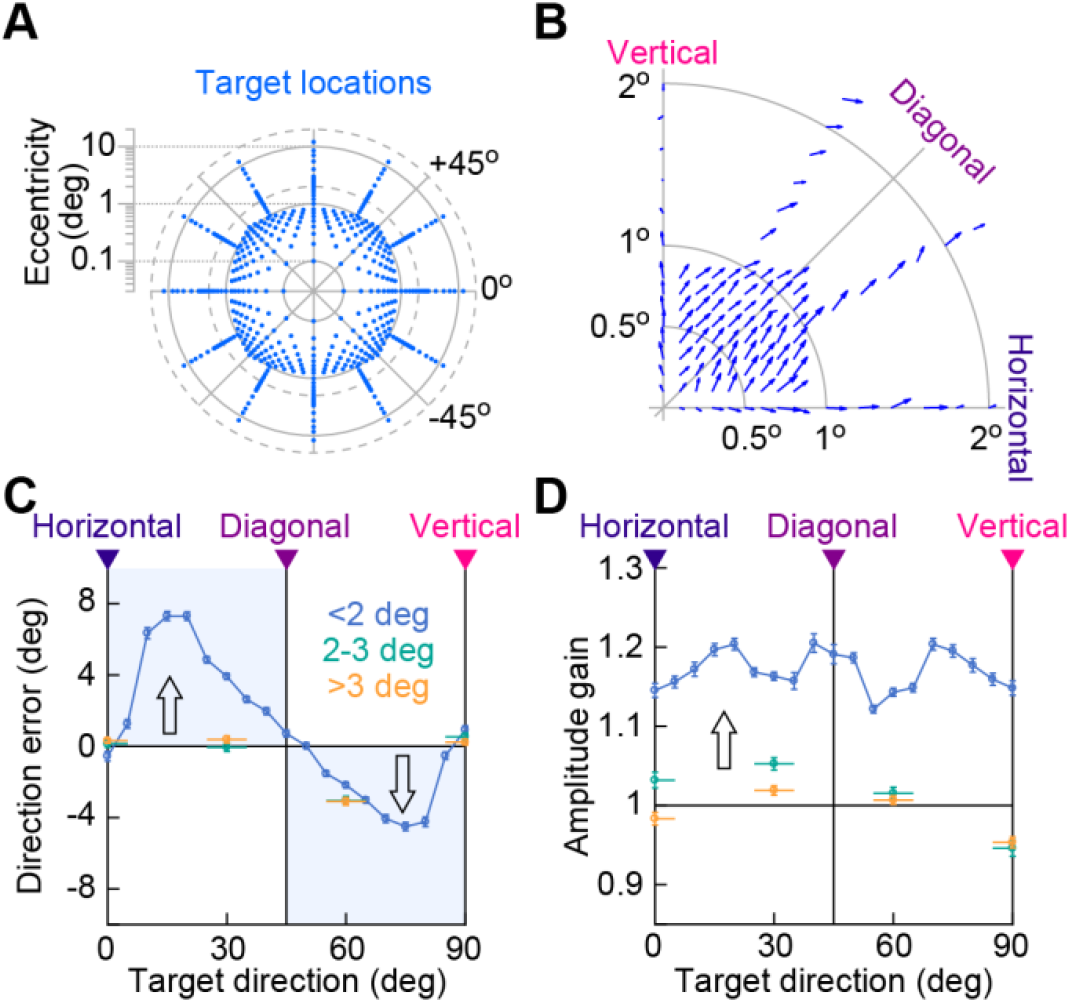
Similar distortion of foveal visual location memory with a different response modality and a foveal visual landmark during the response phase. **(A)** Target locations from our mouse pointing task. The trials were identical, except that during the response phase, the central fixation spot remained visible, and subjects were to maintain gaze on it. Also, the subjects responded with a mouse click at the remembered target location. **(B)** A similar diagonal bias as in the original experiment (Fig. 2) can be seen. **(C, D)** Analyses similar to those of Fig. 3 showing similar results to the original experiment, but the errors were generally smaller in the current experiment. Error bars denote s.e.m. across all responses. Figure S2E-H shows the influence of memory interval length on the response distortions, and Fig. S4 shows results from individual subjects.

We replicated all of the results from the first experiment. Specifically, even though the localization errors were smaller, in general, we still found a diagonal bias for remembered foveal visual eccentricities (Fig. 4B, C). This bias was also significantly weaker for extrafoveal locations (Fig. 4C), again consistent with Figs. 1-3. Similarly, foveal visual eccentricities were associated with substantial overshoots of target locations, which were absent, and sometimes replaced by undershoots (known foveal biases), at extrafoveal eccentricities (Fig. 4D). Moreover, when we evaluated the influence of memory interval duration like we did above, we again found an increased overshoot distortion with longer memory intervals as in the first experiment (Fig. S2E-G), and this effect was not explained by altered reaction times (Fig. S2H). Finally, all of these results were robust at the individual subject level (Fig. S4): all subjects showed a diagonal direction bias in the fovea, and all but one subject showed amplitude overshoots. Importantly, 4 of 7 subjects who performed this version of the task also performed the earlier one, and Figs. S1, S4 highlight one subject’s results in both variants of the experiment (with consistent observations).

Therefore, all of the primary results from Figs. 1-3 were replicated with an altered response modality in which we maintained retinotopic correspondence during stimulus presentation and response execution, and when we also maintained visual correspondence by keeping the foveal visual landmark persistent on the display during both phases of the task.

### Qualitatively different memory-based foveal distortions when reporting target locations with saccades

We finally asked whether distortion of the foveal visual image region during our memory-guided localization task was specific to manual responses. We repeated the same mouse pointing experiment (Fig. 4A), but this time while removing the fixation spot at trial end (and not showing any mouse pointer). Subjects were instructed to make a saccade, including microsaccades (Hafed & Goffart, 2020; Willeke et al., 2019), towards the remembered target location, making this task the same as that used in a large number of classic short term memory studies. The subjects readily made the memory-guided saccades, even for small foveal eccentricities, as we recently described (Hafed & Goffart, 2020; Willeke et al., 2019). Moreover, their reaction times were much faster than in the manual versions of the task (Fig. S3C). So, we asked whether their endpoints also revealed strong distortions of the foveal visual image region during oculomotor-based memory localization.

Figure 5A shows the mislocalizations that we observed. It was immediately evident that the foveal visual image region was also severely distorted. However, there was an upward bias in small saccade endpoints, which would have been masked by remapping all quadrants into a single quadrant (like we had done above). Therefore, in Fig. 5A, we remapped the target and response locations into one hemifield, rather than into one quadrant, in order to maintain representation of lower visual field target locations. Foveal targets in the lower visual field (like upper visual field foveal targets) were now associated with an upward bias in small eye movement endpoints. Therefore, oculomotor-based reporting of target locations from short term memory was still associated with systematic distortions of the foveal visual image region, but the nature of the distortions was qualitatively very different.

**Figure 5.**
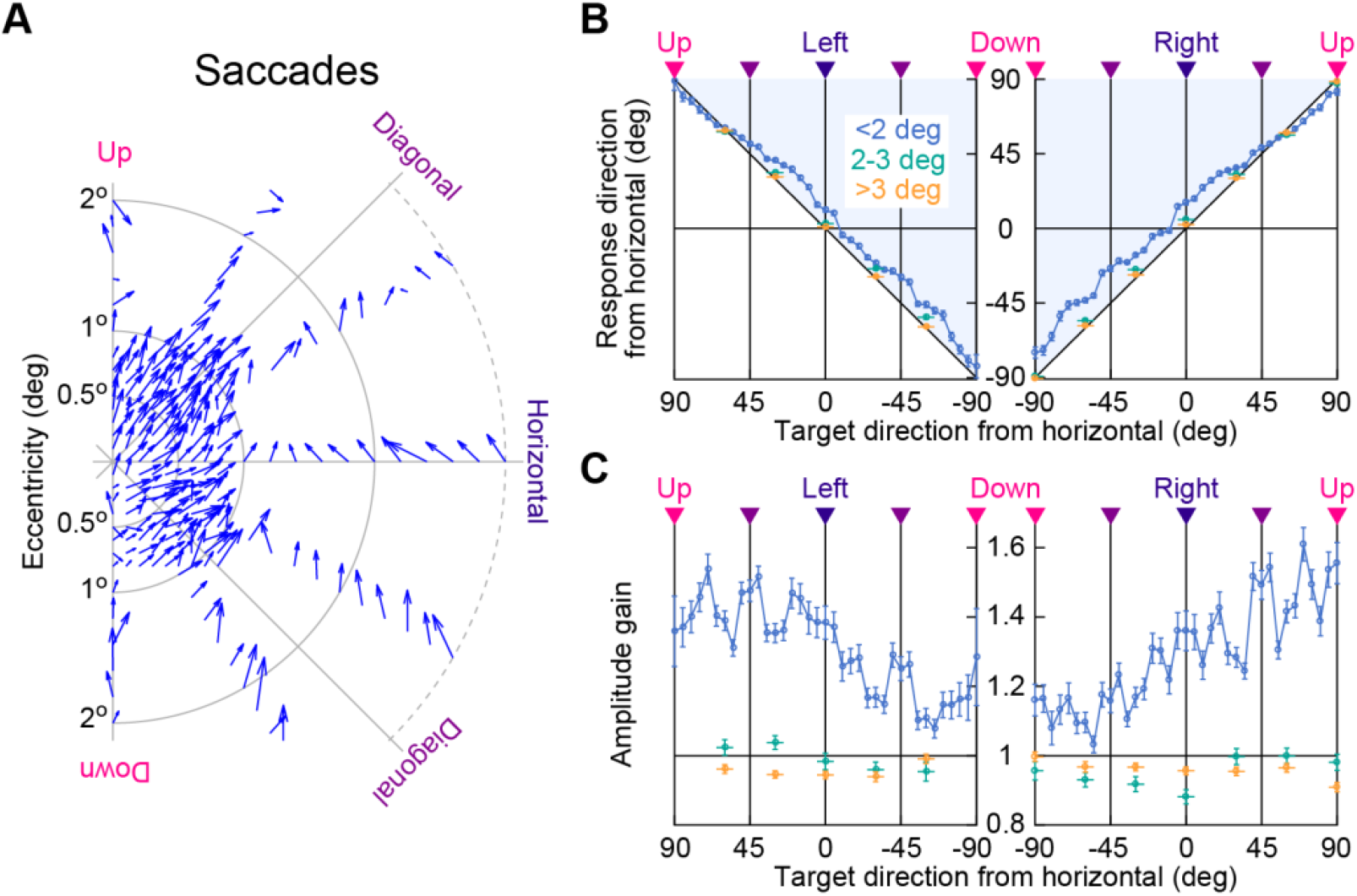
Indicating remembered foveal visual target locations by small saccades is also strongly distorted, but in a qualitatively different manner. **(A)** We repeated the same memory task, but trial end was marked by the removal of the central fixation spot with no other targets; subjects made a saccade towards the remembered target location (which was now blank). We plotted average (foveal) saccade endpoint in relation to true target location, similar to Fig. 2. However, to avoid masking the upward bias that was evident, we now remapped targets and responses into one hemifield instead of one quadrant. Targets in the lower visual field were associated with upward displacements in saccade endpoints, which would have been masked with remapping to a single quadrant. Amplitude overshoots were still evident. **(B)** Response direction from the horizontal meridian as a function of target direction for the right and left visual hemifields. Foveal targets (blue curve) in short term memory were associated with a systematic upward bias (shaded regions). More eccentric targets did not show such a strong bias (green and orange). **(C)** Response amplitude gain for the same target locations. Foveal targets (blue curve) were associated with amplitude overshoots, as in Figs. 1-4. The overshoots got progressively stronger for more upward target locations (consistent with the idea of an overall upward bias for downward targets effectively shortening the intended saccades). Error bars denote s.e.m. across all responses. Figures S5, S6 show individual subject results, as well as demonstration of the qualitative difference in foveal memory distortion between saccadic and manual responses.

To summarize the upward bias that we observed with saccadic memory-based localization, we picked, for all foveal target eccentricities (<2 deg), different target directions from the horizontal meridian, and we plotted the direction of the final response as a function of them (blue curve in Fig. 5B). Response directions were consistently biased upward (shaded regions in Fig. 5B), except for target directions that were near the purely upward meridian: an upward bias for purely upward targets would not bias the response direction, but it would increase the amplitude overshoot. Indeed, in Fig. 5C, we plotted the response amplitude gain for the same data, and we found that the amplitude overshoots associated with remembering foveal visual targets were the largest for the target directions that were near the purely upward direction (blue curve). Thus, oculomotor-based readout of foveal visual target locations from short term memory was still associated with a strong distortion (including amplitude overshoot), but it was dominated by an upward bias rather than a diagonal one.

For extrafoveal target eccentricities, the upward bias was much smaller in amplitude than for foveal targets (green and orange in Fig. 5B), consistent with a significantly milder distortion of extrafoveal eccentricities in our earlier experiments as well (Figs. 1-4). Also consistent with these earlier experiments was the change from amplitude overshoots to amplitude undershoots for extrafoveal targets, which we still observed in the saccade version of the experiment (green and orange in Fig. 5C). Finally, the results in Fig. 5 were robust at the individual subject level (Fig. S5), and we confirmed that within individuals that performed all three variants of the experiments (Methods), the qualitative change in distortion pattern for foveal remembered visual target locations between manual and saccadic responses was still clearly visible (Fig. S6).

## Discussion

We observed a seemingly paradoxical distortion of the highest acuity region of the visual image in short term memory. Attempting to report a remembered foveal visual location after only a few hundred milliseconds from its presentation was associated with significant overshoots in its reported eccentricity, as well as strong biases towards the diagonal axes for non-cardinal target locations. Relative to the original target eccentricities, the distortions in the foveal visual image region were much larger than distortions of extrafoveal eccentricities normally suffering from severe visual acuity losses with sensory stimulation. Oculomotor readout of short term memory for foveal visual target locations was also distorted, but with a qualitatively different pattern of a global upward bias, suggesting that readout modality matters a great deal in studies of visual short term memory.

While the distortions that we observed may be generally related to topographic neural map representations, these distortions are not trivially explained by a simple model of a more diffuse neural trace of the visual target representation in a foveally magnified area, such as the primary visual cortex (Azzopardi & Cowey, 1996; Daniel & Whitteridge, 1961) or the superior colliculus (Chen, Hoffmann, Distler, & Hafed, 2019). For example, readout of a foveally magnified neural map (at different locations) should be similar whether there is a wide hill of activation (diffuse memory trace) versus a narrower hill of activation (which might result from actual stimulus onset). Rather, anisotropies in neural representations of the visual field, that go beyond foveal magnification, might be at play as well (Benson et al., 2021; Silva et al., 2018; Van Essen et al., 1984). Indeed, anisotropies in visual field representations also exist in brain areas relevant for oculomotor readout in our task, as exemplified by the large difference between how the superior colliculus represents the upper and lower visual fields (Hafed & Chen, 2016).

In the visual cortex, anisotropies are related to behavior (Benson et al., 2021; Chapman & Bonhoeffer, 1998; De Valois, Yund, & Hepler, 1982; Li, Peterson, & Freeman, 2003; Silva et al., 2018; Van Essen et al., 1984). For example, visual performance on the horizontal meridian in a variety of tasks is better than performance on the vertical meridian (Carrasco et al., 2001; Talgar & Carrasco, 2002), and also better than performance at oblique locations (Ball & Sekuler, 1987; Heeley & Buchanan-Smith, 1992; Krukowski & Stone, 2005). Besides our diagonal biases, our manual response results with short term memory for foveal target locations demonstrate similar visual meridian patterns (e.g. compare the direction and amplitude errors in Fig. 3 between the horizontal and vertical meridians), and they are in line with evidence that visual meridian performance differences in the periphery persist in short term memory tasks (Montaser-Kouhsari & Carrasco, 2009). In fact, our manual response results can also suggest that anisotropies in representing visual field locations in cortex (Benson et al., 2021; Silva et al., 2018; Van Essen et al., 1984) extend well into the small-eccentricity foveal representations, which makes sense given the large amount of neural tissue magnification associated with foveal eccentricities.

In that regard, it is interesting that differences in perceptual (Appelle, 1972; Ball & Sekuler, 1987; Girshick, Landy, & Simoncelli, 2011; Heeley & Buchanan-Smith, 1992; Krukowski & Stone, 2005; Li et al., 2003; Tomassini, Morgan, & Solomon, 2010) and oculomotor (Krukowski & Stone, 2005) performance often demonstrate a so-called oblique effect, in which performance is worse at oblique directions. In our case, the diagonal bias gave the best directional error performance at oblique directions in the fovea (e.g. Figs. 2, 3A). However, amplitude errors were the largest for the oblique directions (Fig. 3B). Given this diversity of effects, we prefer to use the term diagonal bias, versus oblique effect, as other authors have done (Koehn, Roy, & Barton, 2008) in a different context.

Another reason for our interest in oblique effects is that these earlier studies almost invariably employed a two-interval forced choice paradigm. In such paradigms, a reference stimulus is first presented, followed by a second one. It could be the case that the memory trace of the first stimulus is distorted, as revealed by our results, and that it is this distortion that gives rise to oblique effects in these experiments. As mentioned above, at eccentric locations, visual meridian performance differences indeed occur in a short term memory task (Montaser-Kouhsari & Carrasco, 2009). Moreover, memory distortions can emerge very rapidly, even with memory intervals as short as 50 ms (Werner & Diedrichsen, 2002). Thus, classic oblique effects with 2-interval forced choice paradigms may be mediated by distortions in short term memory like the ones that we have described.

It may also generally be the case that evoking oculomotor or manual reporting from non-visual stimuli would invoke distortions, by virtue of non-visual neural activity that may be needed for successful task performance. For example, in an anti-saccade task, in which subjects generated a saccade to the diametrically opposite location from a visual stimulus, a mild diagonal bias, not unlike ours, was observed for extrafoveal target eccentricities (Koehn et al., 2008). In this case, the distortion emerged with a transformation of the visual signal into a separate movement command, which may or may not require recruitment of short term memory. We even found, in earlier work, repulsion away from the fovea at near eccentricities and towards the fovea at far eccentricities around the time of microsaccade generation (Hafed, 2013), suggesting that any manipulation of neural activity in a logarithmically warped and anisotropic topographic map may be sufficient to cause distortions like the ones that we saw in our study.

Our oculomotor results are also intriguing because they relate to debates on whether saccades in darkness are associated with an upward shift in gaze. This is a robust phenomenon in monkeys (Goffart, Quinet, Chavane, & Masson, 2006; White, Sparks, & Stanford, 1994), but it has been questioned whether humans might also show it (e.g. see related experiments in Goffart et al., 2006; Snodderly, 1987). In our case, complete darkness was not even required to evoke an upward shift in memory-guided saccade endpoints. All that was needed was a saccade to a blank. Importantly, small saccades on the scale of microsaccades were not previously tested for such upward biases under memory guidance. It is possible that it was easier to see upward biases for small saccades exactly because of large foveal magnification, similar to how visual meridian effects (with or without memory) might reflect anisotropies in cortical tissue magnification; our weak extrafoveal effects (relative to monkeys) might, in turn, suggest that humans have larger foveal magnification than monkeys.

In any case, the fact that we still observed upward biases for our humans’ small memory-guided microsaccades is consistent with past observations on the role of active vision at the foveal scale in oculomotor control areas (Hafed, Chen, Tian, Baumann, & Zhang, 2021; Hafed, Yoshida, Tian, Buonocore, & Malevich, 2021). It is also important to remind that saccades (including small ones) are more accurate with visual guidance than without, as we showed recently with the same subjects (and with monkeys) in variants of our memory-based task that had a visual saccade target in the final response phase (Hafed & Goffart, 2020; Willeke et al., 2019).

Our saccade results can also help rule out the possibility that our manual response results were due to a range effect, or the idea that subjects were simply biased towards the midpoint of the range of eccentricities that we tested. Since saccades do not suffer from a range effect (Nuthmann, Vitu, Engbert, & Kliegl, 2016), our observation that saccades still overshot the target at small eccentricities suggest a genuine distortion in memory-based localization, even in manual tasks. This is also consistent with earlier conclusions in extrafoveal memory recall (Kerzel, 2002).

Based on the qualitative difference between our manual response and saccadic results, we also believe that readout modality matters a great deal in studies of short term memory. This is an important point to consider given the large number of studies of short term memory that rely on saccadic responses. Indeed, our results suggest that manual responses in the same task might reveal very different distortions from saccadic responses. Such different distortions may be associated with visual topographic map anisotropies (Fig. 6A), which can be very distinct from oculomotor map anisotropies (Fig. 6B).

**Figure 6.**
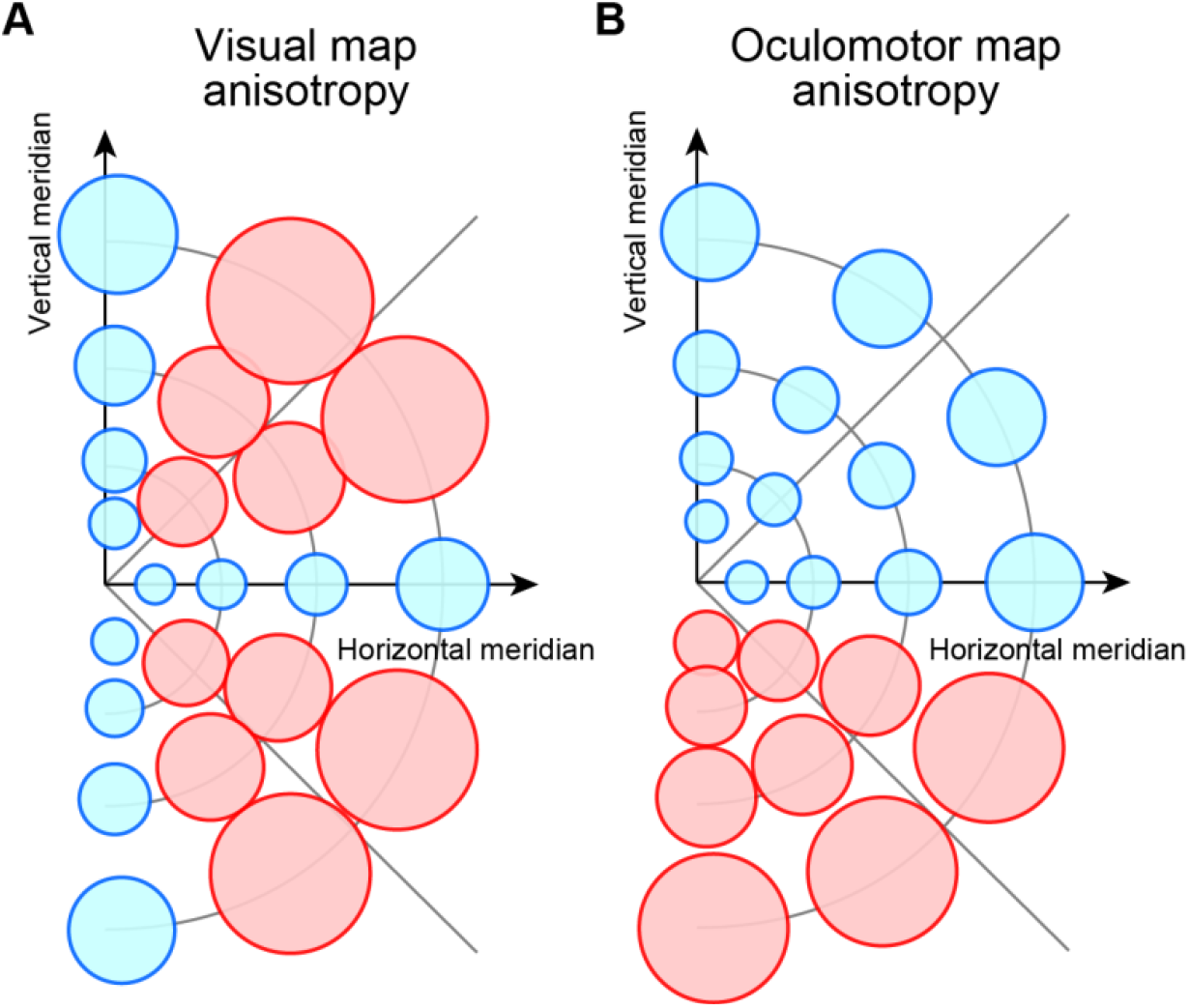
Differences in short term memory distortions across response modalities may be related to differences in topographic anisotropies within the circuits used for readout. **(A)** With manual reporting, anisotropies in cortical visual areas may dominate, with diagonal biases reflecting overrepresentation of the visual meridians (blue) at the expense of non-cardinal locations (red). Here, overrepresentation is schematized as smaller visual receptive fields (circles); note that lower visual field receptive fields are schematized as also being smaller than upper visual field receptive fields. Evidence for such visual field anisotropies exists (Benson et al., 2021; Krukowski & Stone, 2005; Silva et al., 2018; Van Essen et al., 1984) and is consistent with our observations. **(B)** With saccadic reporting, location readout might be more dependent on oculomotor maps, which can have very different anisotropies. For example, there are smaller response fields in the foveal and upper visual field (blue circles) representations of the superior colliculus (Chen et al., 2019; Cynader & Berman, 1972; Hafed & Chen, 2016). This could potentially result in different memory-based distortions with saccadic readout, as we saw in Fig. 5.

Finally, it may also be the case that the fixation spot during the memory interval acted as a visual landmark, which can distort target localization (Schmidt, 2004; Schmidt, Werner, & Diedrichsen, 2003). However, we saw similar distortions with and without a foveal visual landmark during the response phase in our first two experiments. We think that the retinotopic reference frame itself is what induces outward expansion for near locations and foveal biases (Kerzel, 2002; Musseler, van der Heijden, Mahmud, Deubel, & Ertsey, 1999; O’ Regan, 1984; Sheth & Shimojo, 2001) for far locations. To consolidate this idea, it would be interesting in the future to identify foveal visual performance asymmetries (like visual meridian effects) even without a memory task.

In all, our findings highlight the importance of studying multiple aspects of foveal visual processing, particularly at the level of neural mechanisms, since foveal vision, along with its associated active scanning behaviors, is particularly relevant for fully understanding the full spectrum of human visual function.

## Acknowledgements

We were funded by the Werner Reichardt Centre for Integrative Neuroscience (CIN), an excellence cluster (EXC307) funded by the Deutsche Forschungsgemeinschaft (DFG). We were also supported by the DFG-funded Research Unit (FOR1847; project A6; HA6749/2-1).

## Author contributions

All authors conceptualized the experiments. KW and ARC collected the manual response data. ARC and JB collected the saccadic response data. KW and ZMH analyzed the results. ZMH wrote the first draft of the manuscript. All authors edited and finalized the manuscript.

## Methods

We performed novel analyses on behavioral data that were part of another study (Willeke et al., 2019). The earlier study was concerned with memory-guided microsaccades and their associated neural responses in the superior colliculus; the behavioral parts provided necessary control conditions in that study, but they were not analyzed in detail. Here, we analyzed the behavioral parts anew, to reveal properties of foveal visual location recall from short term memory.

### Laboratory setup

The laboratory setup was the same as that used in previous publications (Grujic, Brehm, Gloge, Zhuo, & Hafed, 2018; Hafed, 2013; Willeke et al., 2019). Briefly, subjects were seated 57 cm from the front of a CRT monitor having a refresh rate of 85 Hz, and their heads were stabilized with a custom-built device (Hafed, 2013). We tracked eye movements at 1000 Hz using a video-based eye tracker (EyeLink 1000). The display had a resolution of 41 pixels per deg (Grujic et al., 2018; Idrees, Baumann, Franke, Munch, & Hafed, 2020; Willeke et al., 2019), and it was luminance calibrated. Stimuli used in this study consisted of white dots having 97 cd/m^2^ luminance, which were presented over a uniform gray background of 21 cd/m^2^ luminance (Grujic et al., 2018).

### Subjects

We collected data from a total of 12 subjects. Each variant of the memory experiment was performed by a total of 7 subjects, and 4 subjects participated in all three variants (in separate sessions). In the button cursor movement task, 1 subject was female; in the mouse pointing task, 1 subject was female; and in the memory-guided saccade task, 3 subjects were female. The subjects provided written informed consent prior to their participation, and they were financially compensated for their time.

All experiments were conducted after approval from ethics committees at the Medical Faculty of the University of Tübingen, and the experiments were in line with the Declaration of Helsinki.

The subjects’ ages ranged from 23-39 years.

### Behavioral tasks

#### Button cursor movement task

Each trial started with a central fixation spot (7.3 × 7.3 min arc) presented over a uniform gray background. After a random interval of 750-1250 ms, we briefly flashed a similar spot, for 59 ms, at one of 300 possible locations (indicated in Fig. 1A), spanning eccentricities of 0.1 deg (6 min arc) to 12 deg. 176 out of the 300 target locations had both a horizontal and vertical eccentricity absolute value of less than 1 deg. After a memory period of 300-1100 ms, the fixation spot disappeared, and a response cursor (a crosshair in the shape of a plus sign, having dimensions 59 × 59 min arc) appeared at the center of the display (in place of the initial fixation spot). The subjects pressed one of four buttons on a response box to move the cursor along all four possible cardinal directions, and they then pressed a fifth (middle) button to indicate that the cursor was at the remembered target location. Each button press during cursor movement moved the cursor by 1.46 min arc along a cardinal direction, and prolonged button presses resulted in one of two faster cursor speeds depending on press duration (Willeke et al., 2019). All of this meant that reaction times in this task could be quite long (Fig. S3A), and we allowed subjects up to 20 s for finalizing their response. The subjects were free to move their eyes during cursor movement and final report. After the subjects finalized their response (by pressing the fifth button), the true target location appeared as feedback on how accurate their report was. Each subject participated in 5-6 sessions of 250-350 trials each. We collected a total of 11249 trials across subjects.

#### Mouse pointing task

The task was similar to the one above, except that we kept the central fixation spot visible at trial end (and until subject response). Also, instead of a cursor, the computer mouse pointer became visible to indicate that it was now time to report the remembered target location. The mouse pointer appeared at a random location 1-5 deg in eccentricity away from the true target location. This allowed faster reaction times than in the button cursor movement task, and it also made the subjects’ task somewhat easier, especially since they were now required to maintain their gaze near the central fixation spot at all times while responding (even for the peripheral targets). The subjects’ reporting requirement was to point the mouse pointer at the remembered target location and click the mouse button as quickly as possible. They then received feedback by the reappearance of the target at its original location. As expected, the reaction times in this task were significantly shorter than in the button cursor movement task (Fig. S3B), since moving the mouse pointer could be achieved faster. We used a total of 480 target locations, 296 of which had both a horizontal and a vertical eccentricity of less than 1 deg. Subjects participated in 5 sessions of 380-500 trials each (except for one subject who completed 4 sessions). We collected 16213 trials in this task.

#### Memory-guided saccade task

The task was identical to the mouse pointing task above (including in the target locations). However, at trial end, the fixation spot was removed, and no other stimulus was presented. The removal of the fixation spot was the cue to generate a memory-guided saccade towards the remembered target location. After a prolonged delay well beyond the saccadic reaction times, the target reappeared as feedback to the subjects. We collected 4-5 sessions, of 480-600 trials each, per subject, for a total of 16844 trials.

### Data analyses

Eye movements were analyzed in detail earlier (Hafed & Goffart, 2020; Willeke et al., 2019), including with monkey subjects exhibiting similar effects to the humans (Willeke et al., 2019). Here, we used the already labeled data from these studies to investigate the saccadic distortions shown in Figs. 5, S5, S6.

For manual responses (whether with button cursor movement or mouse pointing), we created tables of reported locations along with true locations. We excluded rare trials in which the subjects missed seeing the flash, for example, due to microsaccadic suppression at flash onset time (Hafed & Krauzlis, 2010). In these cases, subjects typically indicated display center (no cursor movement) or pressed a far location and in the wrong quadrant. Such trials were very few (approximately 0.4%).

We defined amplitude and direction errors as explained in Fig. 1B. Specifically, the average response location across trial repetitions had a vector from display center of a given eccentricity and direction (red in Fig. 1B). The direction error of the response vector was the difference between this vector’s direction from the horizontal meridian and the direction of the vector connecting the display center to the true target location. As for amplitude error, we divided the eccentricity of the average response vector by the eccentricity of the true target location vector. A ratio >1 represented a response amplitude overshoot, and a ratio <1 represented a response amplitude undershoot (e.g. foveal biases for eccentric extrafoveal locations).

For summarizing distortions in foveal location short term memory, we connected the true target location with the average response location using a blue vector (as in the left panel of Fig. 1B). Across all target locations, this resulted in global quiver plots (e.g. Fig. 2A) of distortions. We summarized these distortions in a more compact way by remapping all four quadrants into a representative quadrant, since the manual response tasks revealed symmetry across quadrants (e.g. Fig. 2A). To do this, we reflected the left visual hemifield and its responses to the right hemifield, and we reflected the lower visual field and its responses to the upper visual field. For the saccadic version of the task, it was immediately obvious that the pattern of distortion was not symmetric across all four quadrants (e.g. Fig. 5). Rather, it was symmetric across the right and left visual hemifields only. Therefore, we remapped responses only across the right and left visual hemifields, and we did not reflect the lower visual field into the upper visual field. This would have masked the global upward bias in memory-guided saccade endpoints that we observed (e.g. Fig. 5).

For the manual response tasks, we also created eccentricity and direction bins of the target locations. Because of the sampled locations, the eccentricity bins of 2-3 deg and >3 deg had fewer direction samples than the <2 deg eccentricity bin (e.g. Fig. 1A). Therefore, we did not do further direction binning beyond the categorical direction bins that existed in our sampled target locations. However, for the foveal eccentricities that were critical for this study, we had sufficient direction samples for further direction binning. We created a running average across directions, with direction bin widths of 15 deg and sampled in steps of 5 deg. So, for example, to plot direction error as a function of target direction from the horizontal meridian (e.g. Fig. 3A), we took all target and response reports with eccentricity <2 deg and target direction from the horizontal within a given direction bin. We then plotted the average direction error. We used a similar procedure for amplitude errors. For the analyses of Fig. S2, we also repeated the same procedures but only for trials with either short or long memory intervals. Trials were classified as having short or long memory intervals based on a median split of the time between stimulus flash end (at the beginning of a trial) and trial end (i.e. onset of the response cursor or mouse pointer) across all trials (subsequent reaction times were similar in both groups of trials, as shown in Fig. S2).

For the saccade task, we used similar eccentricity and direction binning. For clarifying the upward bias in direction errors, rather than plotting direction error as a function of direction bin, we plotted the average response direction as a function of direction bin (e.g. Fig. 5B). We also explicitly plotted both right and left visual hemifield results (e.g. Fig. 5B) to demonstrate that the memory-guided saccade direction biases were upward in both hemifields (justifying the hemifield remapping strategy in Fig. 5A).

In all main figures, we plotted results across subjects by pooling their responses together. This was justified because there was a similar number of trials collected per individual subject. However, in the supplementary figures, we also showed all results after first analyzing each subject individually and then averaging the individual subjects’ averages. As expected, the conclusions were the same with either approach.

In all figures, we also provided descriptive statistics with indications of central tendencies and variances around them. We focused on systematic errors in memory recall for our analyses (except for Fig. 1B). This was important for emphasizing that the effects that we reported here could not be attributed to variability in fixational eye position due to fixational eye movements. Specifically, since gaze continuously shifts subliminally during fixation, it may be argued that errors in memory-based target localizations were due to fixational instability. However, if memory-based localization performance was entirely explained by fixational eye movements, then no systematic errors would be observed in our experiments (only non-systematic, or variability, errors would occur). This is because average eye position will be on the fixated target, by definition. For example, in our earlier work (Tian, Yoshida, & Hafed, 2016, 2018), we found that fixational saccades optimize eye position on the fixated target remarkably well, such that average gaze position remains highly accurate.

## Supplementary figures

**Figure S1.**
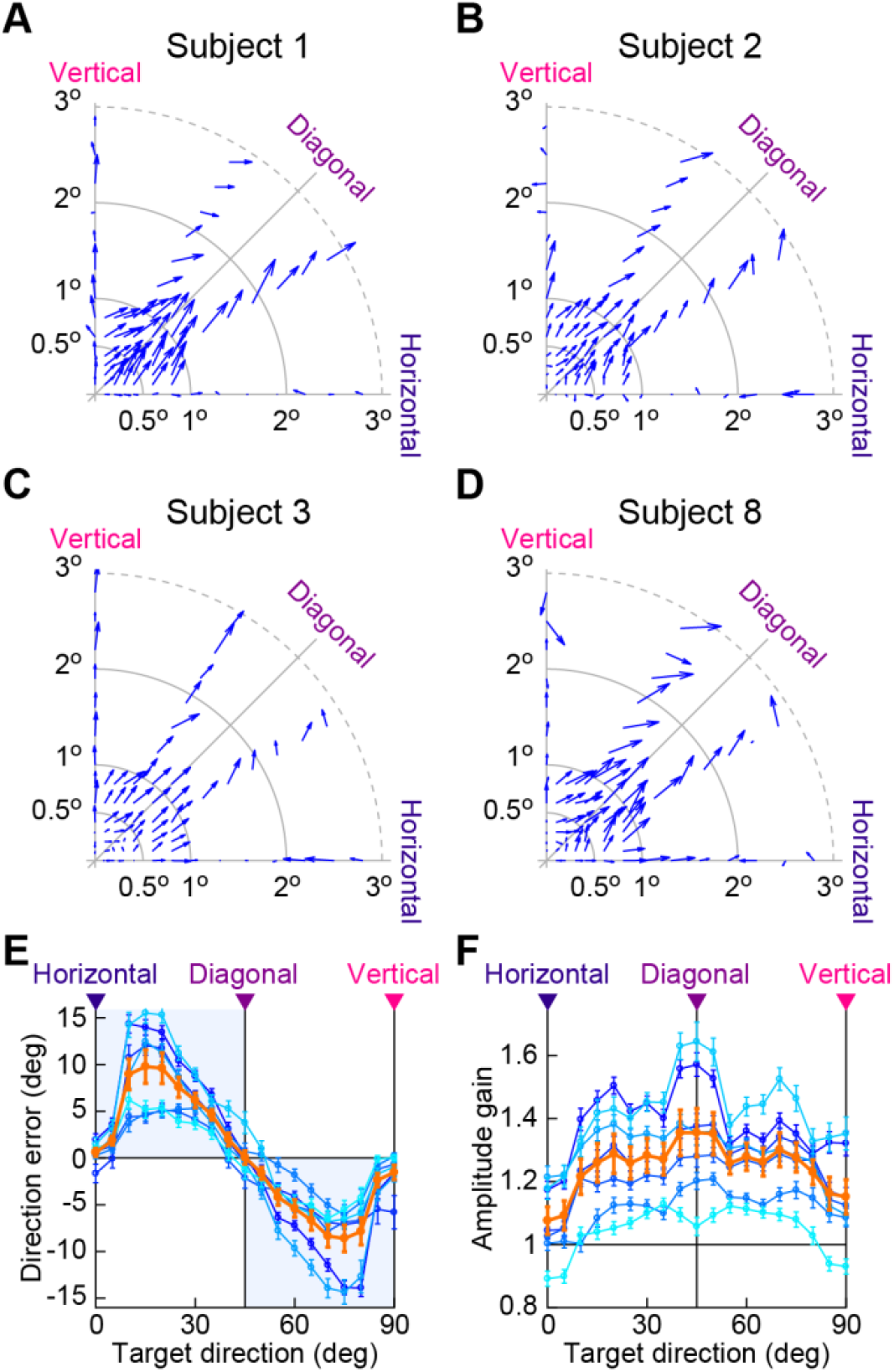
Individual subject results from the experiment of Figs. 1-3. **(A-D)** Four example subjects’ results analyzed in the same way as in Fig. 2B. **(E, F)** All individual subject curves for the analyses of Fig. 3. Each blue-colored curve is a single subject. The thick orange curve shows the average of all individual subject curves, with error bars indicating s.e.m. across the subjects. The error bars in the blue curves indicate s.e.m. across target locations and responses in each eccentricity and direction bin (within subject analysis). For simplicity, we only showed the foveal target eccentricities here, but the results with other eccentricities were all consistent with Figs. 1-3.

**Figure S2.**
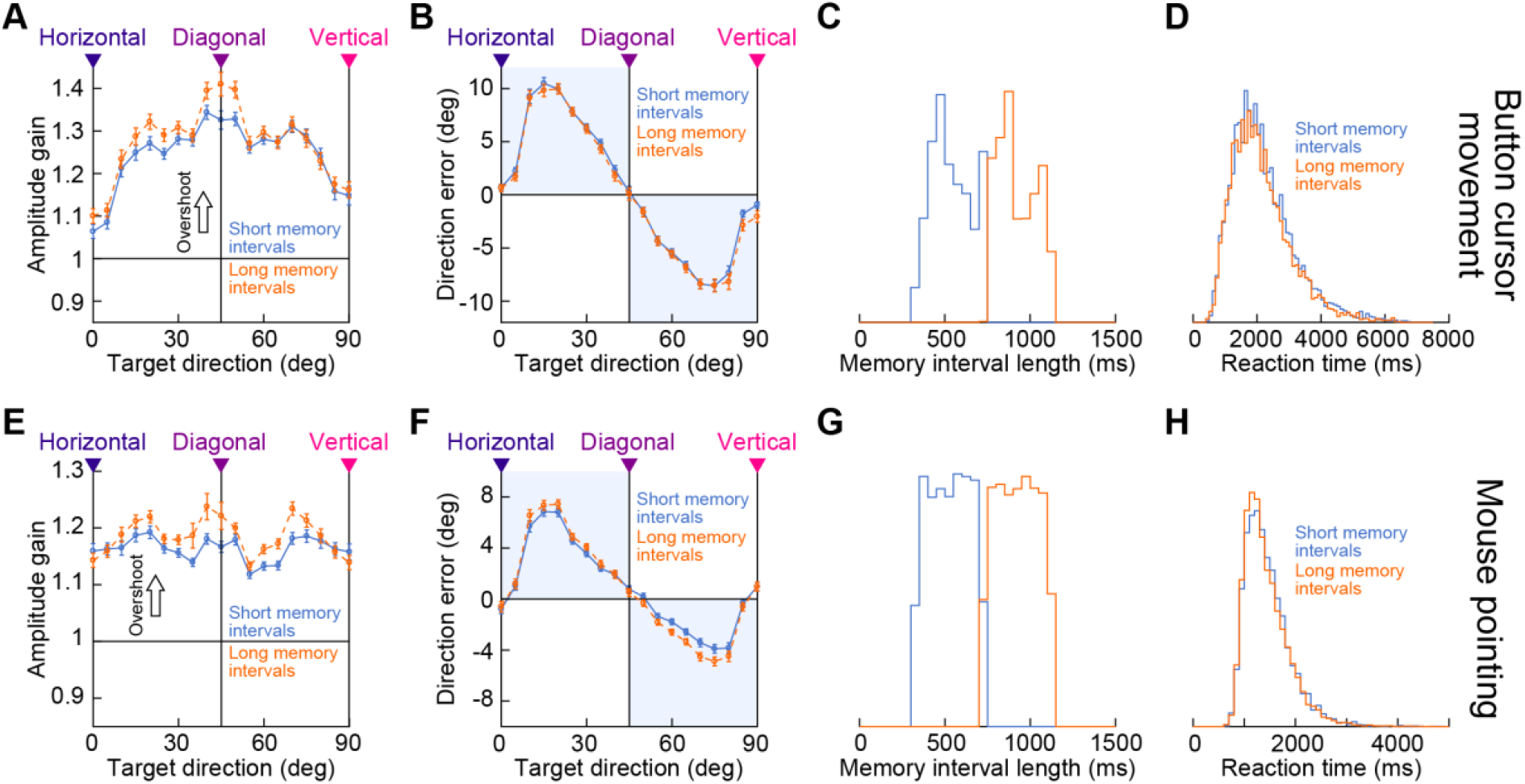
Increased localization distortion with increased memory interval length. **(A)** Response amplitude overshoots for foveal target locations (<2 deg eccentricity) from trials with either a short (solid) or long (dashed) memory interval. Short and long memory intervals were classified based on a median split across all trials (see **C**). Despite the generally short memory intervals used in our study (at most ∼1100 ms), the response amplitude overshoots in the fovea increased in strength for the long memory intervals compared to the short memory intervals. All other conventions are similar to Fig. 3B. **(B)** Direction errors in the fovea (as in Fig. 3A) were less affected by memory interval length, suggesting that memory intervals as short as ∼300 ms were still associated with strong diagonal biases in the foveal visual image region. **(C)** The median split of memory interval length that we used to obtain the results of **A, B. (D)** The increased amplitude overshoots with longer memory intervals in **A** were not explained by different reaction times in the two groups of trials. The distributions of reaction times for the same median split of trials performed in **A**-**C** did not reveal longer reaction times for the long memory interval trials. **(E-H)** Similar results from the mouse pointing task (Fig. 4). Amplitude overshoots increased with longer memory intervals (**E**), and direction errors were also slightly increased (**F**). Finally, like with **D**, the results were largely independent of reaction times across the two groups of trials (**H**). Extra-foveal target locations (not shown here for clarity) had much weaker effects of memory interval length, consistent with the generally weak distortions that they exhibited in general (Figs. 3, 4).

**Figure S3.**
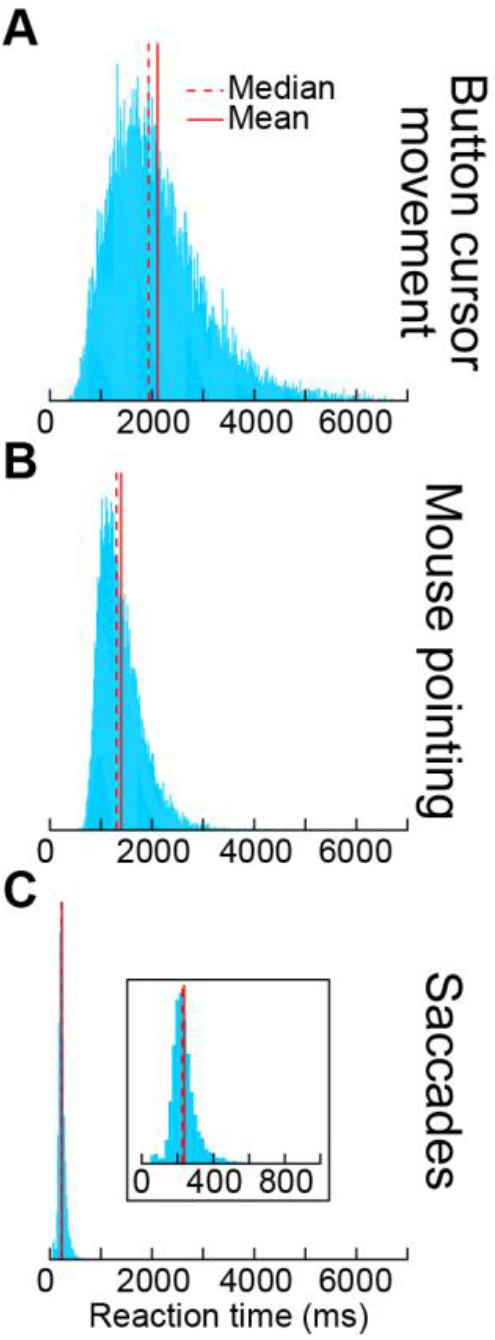
Different time scales of reaction times in the different experiments. **(A)** Reaction time distribution, across all subjects and all trials, from the main experiment of Figs. 1-3 (button cursor movement). The red solid and dashed lines indicate the mean and median reaction times, respectively. **(B)** Reaction time distribution from the mouse pointing experiment. **(C)** With indicating remembered locations by saccades, the reaction times were the shortest (Hafed & Goffart, 2020). The inset magnifies the x-axis for easier visualization of the saccadic reaction times.

**Figure S4.**
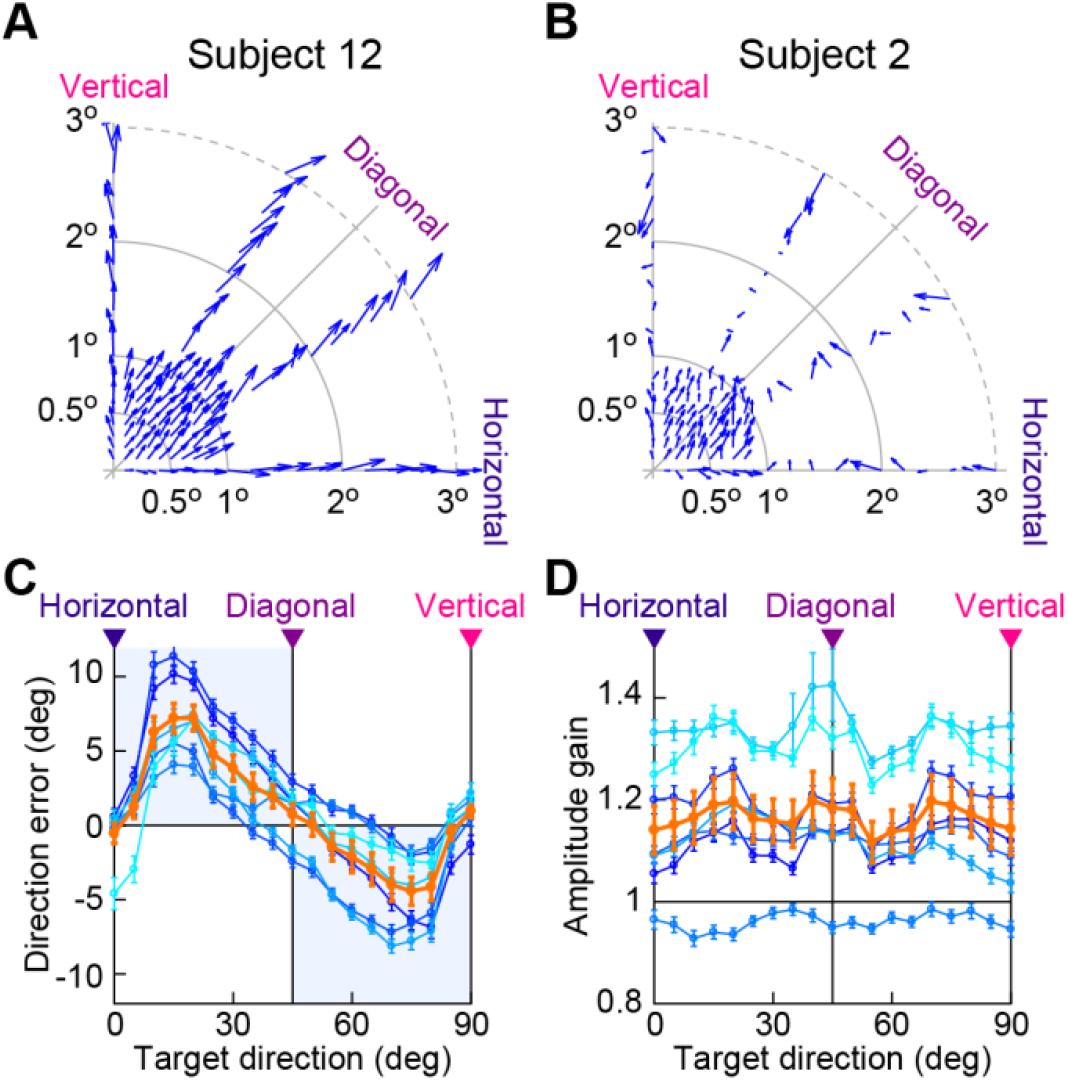
Individual subjects from the experiment of Fig. 4. **(A, B)** Two example subjects demonstrating foveal overshoot and a diagonal bias in the mouse pointing variant of the experiment. Note that **B** shows the same subject as in Fig. S1B, demonstrating an example of an individual who performed both experiments (as well as the saccade variant; see Figs. S5, S6). **(C, D)** Same analyses as in Fig. 4C, D, but now showing individual subject curves. All conventions are the same as in Fig. S1.

**Figure S5.**
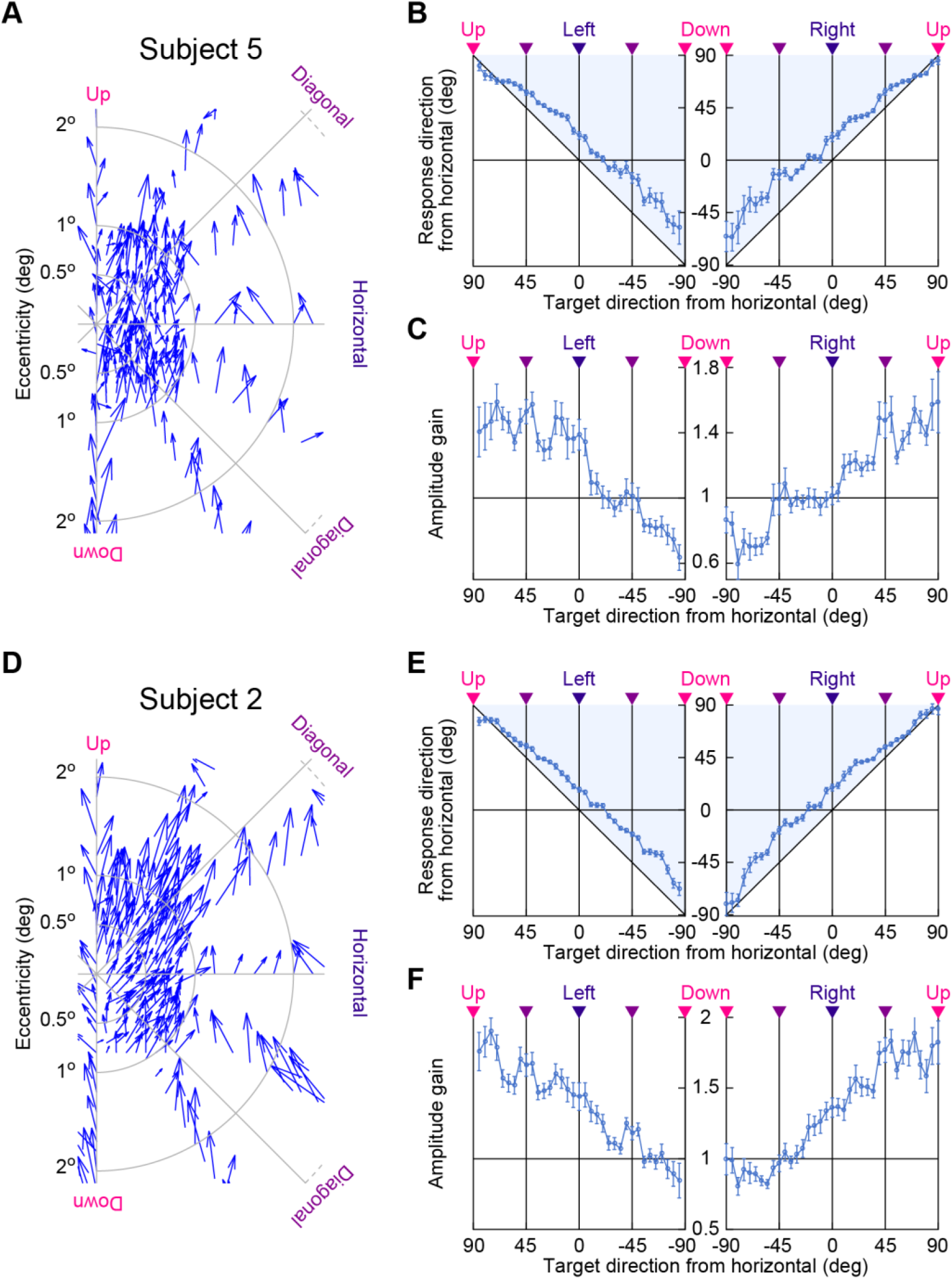
Individual subjects from the oculomotor readout experiment of Fig. 5. **(A)** Quiver plot of target and saccade endpoint locations from one subject. The upward bias of foveal memory-guided saccades that was seen in Fig. 5 is still evident. **(B)** Response directions as a function of target directions for the same subject. This figure is formatted as in Fig. 5B. Only foveal targets are shown for simplicity, and error bars indicate s.e.m. across response locations within a given direction bin (within subject analysis). An upward bias in saccade directions is evident (shaded regions). **(C)** Response amplitude gain as a function of target direction for the same subject. Downward saccades undershot the target slightly (due to an upward bias in endpoint locations), and upward saccades strongly overshot the target. **(D-F)** Same analyses as in **A**-**C**, but for a second example subject. Note that this subject performed all three experiments (see Figs. S1, S4).

**Figure S6.**
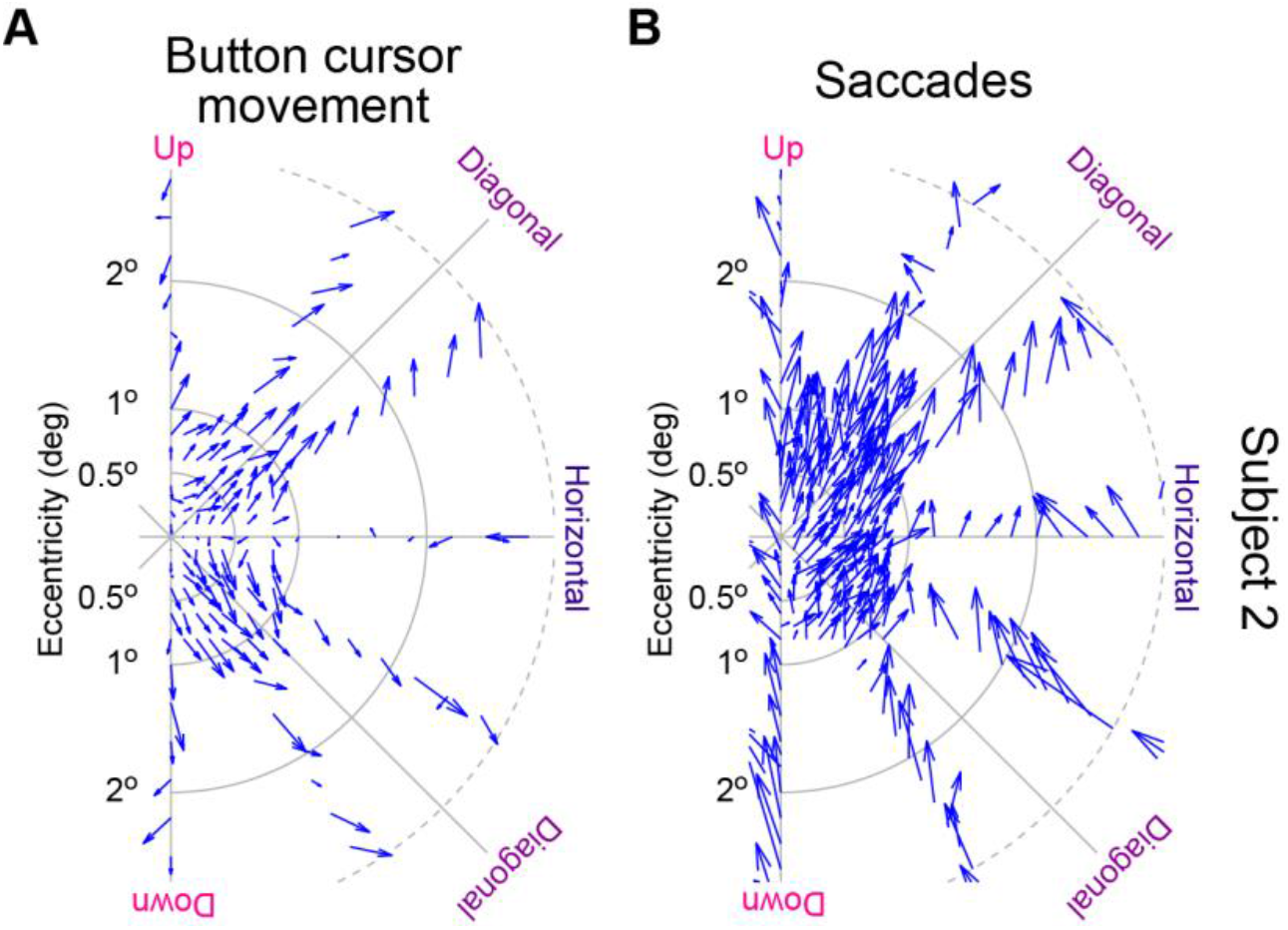
Demonstration of the qualitative difference in foveal location distortion with different response modalities in an individual subject. **(A)** Distortions in the button cursor experiment from one subject who performed all three experiments. The data are now plotted in the same way that we plotted the saccade data. That is, we remapped target and response locations to one hemifield rather than to one quadrant. The subject showed clear overshoots and diagonal biases, as in Figs. 1-3. **(B)** The same subject showed upward biases when asked to indicate the remembered foveal target locations with small saccades.

## References

Appelle, S. (1972). Perception and discrimination as a function of stimulus orientation: the “oblique effect” in man and animals. Psychol Bull, 78(4), 266–278. doi:10.1037/h0033117

Azzopardi, P., & Cowey, A. (1996). The overrepresentation of the fovea and adjacent retina in the striate cortex and dorsal lateral geniculate nucleus of the macaque monkey. Neuroscience, 72(3), 627–639.

Ball, K., & Sekuler, R. (1987). Direction-specific improvement in motion discrimination. Vision Res, 27(6), 953–965. doi:10.1016/0042-6989(87)90011-3

Benson, N. C., Kupers, E. R., Babot, A., Carrasco, M., & Winawer, J. (2021). Cortical Magnification in Human Visual Cortex Parallels Task Performance around the Visual Field. Elife, 10. doi:10.7554/eLife.67685

Bruce, C. J., & Goldberg, M. E. (1985). Primate frontal eye fields. I. Single neurons discharging before saccades. J Neurophysiol, 53(3), 603–635.

Carrasco, M., Talgar, C. P., & Cameron, E. L. (2001). Characterizing visual performance fields: effects of transient covert attention, spatial frequency, eccentricity, task and set size. Spatial vision, 15(1), 61–75.

Chapman, B., & Bonhoeffer, T. (1998). Overrepresentation of horizontal and vertical orientation preferences in developing ferret area 17. Proc Natl Acad Sci U S A, 95(5), 2609–2614. doi:10.1073/pnas.95.5.2609

Chen, C. Y., Hoffmann, K. P., Distler, C., & Hafed, Z. M. (2019). The Foveal Visual Representation of the Primate Superior Colliculus. Curr Biol, 29(13), 2109–2119 e2107. doi:10.1016/j.cub.2019.05.040

Constantinidis, C., & Procyk, E. (2004). The primate working memory networks. Cogn Affect Behav Neurosci, 4(4), 444–465. doi:10.3758/cabn.4.4.444

Cynader, M., & Berman, N. (1972). Receptive-field organization of monkey superior colliculus. J Neurophysiol, 35(2), 187–201.

Daniel, P. M., & Whitteridge, D. (1961). The representation of the visual field on the cerebral cortex in monkeys. J Physiol, 159, 203–221.

De Valois, R. L., Yund, E. W., & Hepler, N. (1982). The orientation and direction selectivity of cells in macaque visual cortex. Vision Res, 22(5), 531–544. doi:10.1016/0042-6989(82)90112-2

Edelman, J. A., & Goldberg, M. E. (2001). Dependence of saccade-related activity in the primate superior colliculus on visual target presence. J Neurophysiol, 86(2), 676–691. doi:10.1152/jn.2001.86.2.676

Funahashi, S., Bruce, C. J., & Goldman-Rakic, P. S. (1989). Mnemonic coding of visual space in the monkey’ s dorsolateral prefrontal cortex. J Neurophysiol, 61(2), 331–349. doi:10.1152/jn.1989.61.2.331

Girshick, A. R., Landy, M. S., & Simoncelli, E. P. (2011). Cardinal rules: visual orientation perception reflects knowledge of environmental statistics. Nat Neurosci, 14(7), 926–932. doi:10.1038/nn.2831

Gnadt, J. W., & Andersen, R. A. (1988). Memory related motor planning activity in posterior parietal cortex of macaque. Exp Brain Res, 70(1), 216–220. doi:10.1007/BF00271862

Gnadt, J. W., Bracewell, R. M., & Andersen, R. A. (1991). Sensorimotor transformation during eye movements to remembered visual targets. Vision Res, 31(4), 693–715. doi:10.1016/0042-6989(91)90010-3

Goffart, L., Quinet, J., Chavane, F., & Masson, G. S. (2006). Influence of background illumination on fixation and visually guided saccades in the rhesus monkey. Vision Res, 46(1-2), 149-162. doi:10.1016/j.visres.2005.07.026

Grujic, N., Brehm, N., Gloge, C., Zhuo, W., & Hafed, Z. M. (2018). Peri-saccadic perceptual mislocalization is different for upward saccades. J Neurophysiol. doi:10.1152/jn.00350.2018

Hafed, Z. M. (2013). Alteration of visual perception prior to microsaccades. Neuron, 77(4), 775–786. doi:10.1016/j.neuron.2012.12.014

Hafed, Z. M., & Chen, C. Y. (2016). Sharper, Stronger, Faster Upper Visual Field Representation in Primate Superior Colliculus. Curr Biol, 26(13), 1647–1658. doi:10.1016/j.cub.2016.04.059

Hafed, Z. M., Chen, C. Y., Tian, X., Baumann, M., & Zhang, T. (2021). Active vision at the foveal scale in the primate superior colliculus. J Neurophysiol. doi:10.1152/jn.00724.2020

Hafed, Z. M., & Goffart, L. (2020). Gaze direction as equilibrium: more evidence from spatial and temporal aspects of small-saccade triggering in the rhesus macaque monkey. J Neurophysiol, 123(1), 308–322. doi:10.1152/jn.00588.2019

Hafed, Z. M., & Krauzlis, R. J. (2010). Microsaccadic suppression of visual bursts in the primate superior colliculus. J Neurosci, 30(28), 9542–9547. doi:30/28/9542 [pii] 10.1523/JNEUROSCI.1137-10.2010

Hafed, Z. M., Yoshida, M., Tian, X., Buonocore, A., & Malevich, T. (2021). Dissociable Cortical and Subcortical Mechanisms for Mediating the Influences of Visual Cues on Microsaccadic Eye Movements. Front Neural Circuits, 15, 638429. doi:10.3389/fncir.2021.638429

Heeley, D. W., & Buchanan-Smith, H. M. (1992). Directional acuity for drifting plaids. Vision Res, 32(1), 97–104. doi:10.1016/0042-6989(92)90117-2

Hikosaka, O., & Wurtz, R. H. (1983). Visual and oculomotor functions of monkey substantia nigra pars reticulata. III. Memory-contingent visual and saccade responses. J Neurophysiol, 49(5), 1268–1284.

Idrees, S., Baumann, M. P., Franke, F., Munch, T. A., & Hafed, Z. M. (2020). Perceptual saccadic suppression starts in the retina. Nature communications, 11(1), 1977. doi:10.1038/s41467-020-15890-w

Kerzel, D. (2002). Memory for the position of stationary objects: disentangling foveal bias and memory averaging. Vision Res, 42(2), 159–167.

Koehn, J. D., Roy, E., & Barton, J. J. (2008). The “diagonal effect”: a systematic error in oblique antisaccades. J Neurophysiol, 100(2), 587–597. doi:10.1152/jn.90268.2008

Kojima, S. (1980). Prefrontal unit activity in the monkey: relation to visual stimuli and movements. Exp Neurol, 69(1), 110–123. doi:10.1016/0014-4886(80)90147-8

Krukowski, A. E., & Stone, L. S. (2005). Expansion of direction space around the cardinal axes revealed by smooth pursuit eye movements. Neuron, 45(2), 315–323. doi:10.1016/j.neuron.2005.01.005

Leavitt, M. L., Pieper, F., Sachs, A. J., & Martinez-Trujillo, J. C. (2018). A Quadrantic Bias in Prefrontal Representation of Visual-Mnemonic Space. Cereb Cortex, 28(7), 2405–2421. doi:10.1093/cercor/bhx142

Li, B., Peterson, M. R., & Freeman, R. D. (2003). Oblique effect: a neural basis in the visual cortex. J Neurophysiol, 90(1), 204–217. doi:10.1152/jn.00954.2002

Liu, T., Heeger, D. J., & Carrasco, M. (2006). Neural correlates of the visual vertical meridian asymmetry. Journal of vision, 6(11), 1294–1306. doi:10.1167/6.11.12

Mays, L. E., & Sparks, D. L. (1980). Dissociation of visual and saccade-related responses in superior colliculus neurons. J Neurophysiol, 43(1), 207–232.

Mohler, C. W., & Wurtz, R. H. (1976). Organization of monkey superior colliculus: intermediate layer cells discharging before eye movements. J Neurophysiol, 39(4), 722–744.

Montaser-Kouhsari, L., & Carrasco, M. (2009). Perceptual asymmetries are preserved in short-term memory tasks. Attention, perception & psychophysics, 71(8), 1782–1792. doi:10.3758/APP.71.8.1782

Musseler, J., van der Heijden, A. H., Mahmud, S. H., Deubel, H., & Ertsey, S. (1999). Relative mislocalization of briefly presented stimuli in the retinal periphery. Percept Psychophys, 61(8), 1646–1661. doi:10.3758/bf03213124

Nuthmann, A., Vitu, F., Engbert, R., & Kliegl, R. (2016). No Evidence for a Saccadic Range Effect for Visually Guided and Memory-Guided Saccades in Simple Saccade-Targeting Tasks. PLoS One, 11(9), e0162449. doi:10.1371/journal.pone.0162449

O’Regan, J. K. (1984). Retinal versus extraretinal influences in flash localization during saccadic eye movements in the presence of a visible background. Percept Psychophys, 36(1), 1–14.

Schmidt, T. (2004). Spatial distortions in visual short-term memory: Interplay of intrinsic and extrinsic reference systems. Spatial Cognition and Computation, 4(4), 313–336. doi:10.1207/s15427633scc0404_2

Schmidt, T., Werner, S., & Diedrichsen, J. (2003). Spatial distortions induced by multiple visual landmarks: how local distortions combine to produce complex distortion patterns. Percept Psychophys, 65(6), 861–873. doi:10.3758/bf03194820

Sheth, B. R., & Shimojo, S. (2001). Compression of space in visual memory. Vision Res, 41(3), 329–341.

Silva, M. F., Brascamp, J. W., Ferreira, S., Castelo-Branco, M., Dumoulin, S. O., & Harvey, B. M. (2018). Radial asymmetries in population receptive field size and cortical magnification factor in early visual cortex. Neuroimage, 167, 41–52. doi:10.1016/j.neuroimage.2017.11.021

Snodderly, D. M. (1987). Effects of light and dark environments on macaque and human fixational eye movements. Vision Res, 27(3), 401–415. doi:0042-6989(87)90089-7 [pii]

Talgar, C. P., & Carrasco, M. (2002). Vertical meridian asymmetry in spatial resolution: visual and attentional factors. Psychonomic bulletin & review, 9(4), 714–722.

Tian, X., Yoshida, M., & Hafed, Z. M. (2016). A Microsaccadic Account of Attentional Capture and Inhibition of Return in Posner Cueing. Front Syst Neurosci, 10, 23. doi:10.3389/fnsys.2016.00023

Tian, X., Yoshida, M., & Hafed, Z. M. (2018). Dynamics of fixational eye position and microsaccades during spatial cueing: the case of express microsaccades. J Neurophysiol, 119(5), 1962–1980. doi:10.1152/jn.00752.2017

Tomassini, A., Morgan, M. J., & Solomon, J. A. (2010). Orientation uncertainty reduces perceived obliquity. Vision Res, 50(5), 541–547. doi:10.1016/j.visres.2009.12.005

Van Essen, D. C., Newsome, W. T., & Maunsell, J. H. (1984). The visual field representation in striate cortex of the macaque monkey: asymmetries, anisotropies, and individual variability. Vision Res, 24(5), 429–448.

Werner, S., & Diedrichsen, J. (2002). The time course of spatial memory distortions. Mem Cognit, 30(5), 718–730. doi:10.3758/bf03196428

White, J. M., Sparks, D. L., & Stanford, T. R. (1994). Saccades to remembered target locations: an analysis of systematic and variable errors. Vision Res, 34(1), 79–92. doi:10.1016/0042-6989(94)90259-3

Willeke, K. F., Tian, X., Buonocore, A., Bellet, J., Ramirez-Cardenas, A., & Hafed, Z. M. (2019). Memory-guided microsaccades. Nature communications, 10(1), 3710. doi:10.1038/s41467-019-11711-x

Wimmer, K., Nykamp, D. Q., Constantinidis, C., & Compte, A. (2014). Bump attractor dynamics in prefrontal cortex explains behavioral precision in spatial working memory. Nat Neurosci, 17(3), 431–439. doi:10.1038/nn.3645

